# Mitochondrial hypoxic stress induces widespread RNA editing by APOBEC3G in lymphocyte

**DOI:** 10.1101/389791

**Authors:** Shraddha Sharma, Jianmin Wang, Scott Portwood, Eduardo Cortes-Gomez, Orla Maguire, Per H. Basse, Eunice S. Wang, Bora E. Baysal

**Affiliations:** Department of Pathology and Laboratory Medicine; Department of Bioinformatics and Biostatistics; Department of Medicine; Department of Flow and Image Cytometry; Roswell Park Comprehensive Cancer Center, Buffalo, NY 14263, USA

**Keywords:** RNA editing, APOBEC3, NK cells, hypoxia, cell stress, mitochondria, epitranscriptome, innate immune cells

## Abstract

Protein recoding by RNA editing is required for normal health and evolutionary adaptation. However, *de novo* induction of RNA editing in response to environmental factors is an uncommon phenomenon. While APOBEC3A edits many mRNAs in monocytes/macrophages in response to hypoxia and interferons, the physiological significance of such editing is unclear. Here we show that the related APOBEC3G cytidine deaminase induces site-specific C-to-U RNA editing in natural killer (NK), CD8+ T cells and lymphoma cell lines upon cellular crowding and hypoxia. RNASeq analysis of hypoxic NK cells reveals widespread C-to-U recoding mRNA editing that is enriched for genes involved in mRNA translation. APOBEC3G promotes Warburg-like metabolic remodeling and reduces proliferation of HuT78 T cells under similar conditions. Hypoxia-induced RNA editing by APOBEC3G can be mimicked by the inhibition of mitochondrial respiration, and occurs independently of HIF-1α. Thus, APOBEC3G is an endogenous RNA editing enzyme, which is induced by mitochondrial hypoxic stress to promote adaptation in lymphocytes.

## Background

RNA editing is an evolutionarily conserved post-transcriptional modification that can result in amino acid recoding and altered protein function [1]. Protein recoding RNA editing plays an important role during development and in helping organisms adapt to changes in the environment. A-to-I (A>I) and C-to-U (C>U) are the two most common types of RNA editing in mammals, carried out by the ADAR and APOBEC enzymes, respectively.

The more widespread ADAR-mediated A>I RNA editing, mostly occurs (~98%) in non-coding repetitive regions [2], likely to combat viral infection and to regulate innate immunity; to prevent retrotransposon insertion in the genome [3]; or to affect the RNA processing pathway [4]. Environmental factors such as hypoxia or neural activity can modify the level of A>I editing in RNAs of certain genes[5, 6], which are already edited under normal physiological conditions (baseline). Recent studies suggest that the evolutionary acquisition of A>I RNA editing sites can facilitate temperature adaptation in octopus, flies, and single cell organisms [7-10]. However, whether or not RNA editing can be dynamically induced at specific sites *de novo* in response to environmental factors, especially in mammals, is not understood well.

In mammals, C>U RNA editing by cytidine deamination is infrequent in baseline transcript sequences under normal physiological conditions. An exception is APOBEC1-mediated RNA editing, which is mainly involved in the production of short isoform of the ApoB protein in intestinal cells [11]. The related APOBEC3 (A3) family of enzymes [12, 13], consisting of A3A, -B, -C, -D, -F, -G and -H are widely considered as antiviral innate restriction factors because they can mutate foreign genetic material (mainly ssDNA) and inhibit their replication in *in vitro* models [14]. Recently, we described that APOBEC3A (A3A) induces widespread RNA editing resulting in protein recoding of dozens of genes in primary monocytes when cultured at a high density under hypoxia (low oxygen) or when exposed to interferons (IFN-1 and IFN-γ), and in M1 type macrophages as a result of IFN-γ treatment [15]. However, the relationship between viral restriction and cellular RNA editing by A3A, and the functional significance of such editing is unknown.

APOBEC3G (A3G), the most studied member of the A3 family, incorporates into vif-deficient HIV-1 virions, and inhibits HIV-1 replication in target cells by causing crippling C>U mutations in its minus ssDNA strand and by inhibiting reverse transcription [14, 16]. Interestingly, we found that exogenous transient expression of A3G in HEK293T cells causes C>U editing in mRNAs of hundreds of genes, which are largely distinct from those edited by A3A [17]. While these findings indicate that the A3G enzyme is capable of RNA editing, whether or not such editing occurs in primary cells under physiologically relevant conditions is unknown. Therefore, we hypothesized that A3G-mediated RNA editing will be induced in cells which express this enzyme.

In this study, we analyze the cell type specific expression of A3G and identify widespread RNA editing mediated by A3G, induced by high cell density and hypoxia in NK, CD8+ T cells and in the widely studied Hut78 T cell line. Our findings reveal that under hypoxic stress, A3G-mediated RNA editing converges at targets involved in mRNA translation, likely to reorganize the cellular translation apparatus. Furthermore, we show that A3G promotes adaptation to hypoxic stress by suppressing cell proliferation and by promoting glycolysis over mitochondrial respiration. Thus, A3G is a novel endogenous RNA editing enzyme which can facilitate cellular adaptation to mitochondrial hypoxic cell stress in cytotoxic lymphocytes.

## Results

### Cell type specific expression of APOBEC3G

To examine the endogenous RNA editing activity of A3G, we first analyzed A3G’s cell type specific expression levels. Since RNA editing by A3A is observed in cell types that highly express A3A (monocytes and macrophages), we reasoned that A3G-mediated RNA editing would be more likely to occur in cells that highly express this enzyme. A meta-analysis of the publicly available microarray datasets [18] indicated high expression of A3G in gamma delta T cells, NK cells and CD8+ T cells (in that order, Fig. 1a). Individual gene expression datasets including GeneAtlas U133A [19] and immune-response *in silico* (IRIS) [20] confirmed a higher expression of A3G in NK cells relative to T, B lymphocytes and myeloid cells. We experimentally confirmed high expression levels of A3G in primary NK and CD8+ T cells, but found lower expression in primary CD4+ T cells purified from peripheral blood (Fig. 1b). These results are unexpected because prior studies have implied a potential functional role of A3G in restricting HIV-1 in infected CD4+ T cells [21, 22] whereas other studies did not include NK cells in APOBEC3 gene expression profiling [23, 24]. In contrast, our findings reveal the highest expression of A3G in NK and CD8+T lymphocytes that are not infected by HIV-1.

### Identification of RNA editing by APOBEC3G in NK cells

We have previously shown that A3A, which is highly expressed in monocytes and macrophages shows very low or the absence of RNA editing in these cells when freshly isolated from peripheral blood mononuclear cells (PBMCs) [15]. However, RNA editing is induced when monocytes/macrophages are cultured at a high cell density and low oxygen (hypoxia, 1% O_2_) or by interferons [15, 25]. Since A3G is highly expressed in NK cells, we hypothesized that RNA editing will be induced in NK cells when subjected to hypoxia and/or high cell density. We cultured PBMCs for 40 hours at a high cell density (5×10^7^ cells in 1.8 ml per well in a 12-well plate) under normoxia or hypoxia and isolated NK cells. Under these conditions, we observed upregulation of the phosphorylated α subunit of the eukaryotic initiation factor-2 (eIF-2α) at Ser 51-a conserved event activated in response to various cell stresses including hypoxia [26] at 20 h, suggesting that NK cells were stressed (Fig. 1c). To examine site-specific C>U editing in RNAs of NK cells, we selected several candidate genes including *TM7SF3* that we have previously shown high-level RNA editing on overexpressing A3G in 293T cells [17]. *TM7SF3* did not show any RNA editing in freshly isolated (T0/baseline) NK cells (Fig. 1d). However, we found evidence for induction of RNA editing in *TM7SF3* due to cellular crowding with/without hypoxia (higher in hypoxia) (Fig. 1d), which did not further increase with IFN-γ treatment (Additional file 1; Figure S1a). Since A3G is also expressed in CD8+ T cells and to a lesser extent in CD4+ T cells (Fig. 1a), we cultured PBMCs as mentioned above and isolated NK, CD8+ and CD4+ cell subsets from the same donors. Site-specific RNA editing (>5%) was observed in NK cells and to a lesser extent in CD8+ T cells, but not in CD4+ T cells (Fig. 1e), in parallel with the relative expression levels of A3G in these cell types. Since editing in NK and CD8+ T cells occurs in RNAs of genes that have been previously shown to be edited in the 293T/A3G overexpression system (*TM7SF3, RPL10A, RFX7*), our results suggest that A3G induces RNA editing in cytotoxic lymphocytes, particularly in NK cells.

**Fig. 1.**
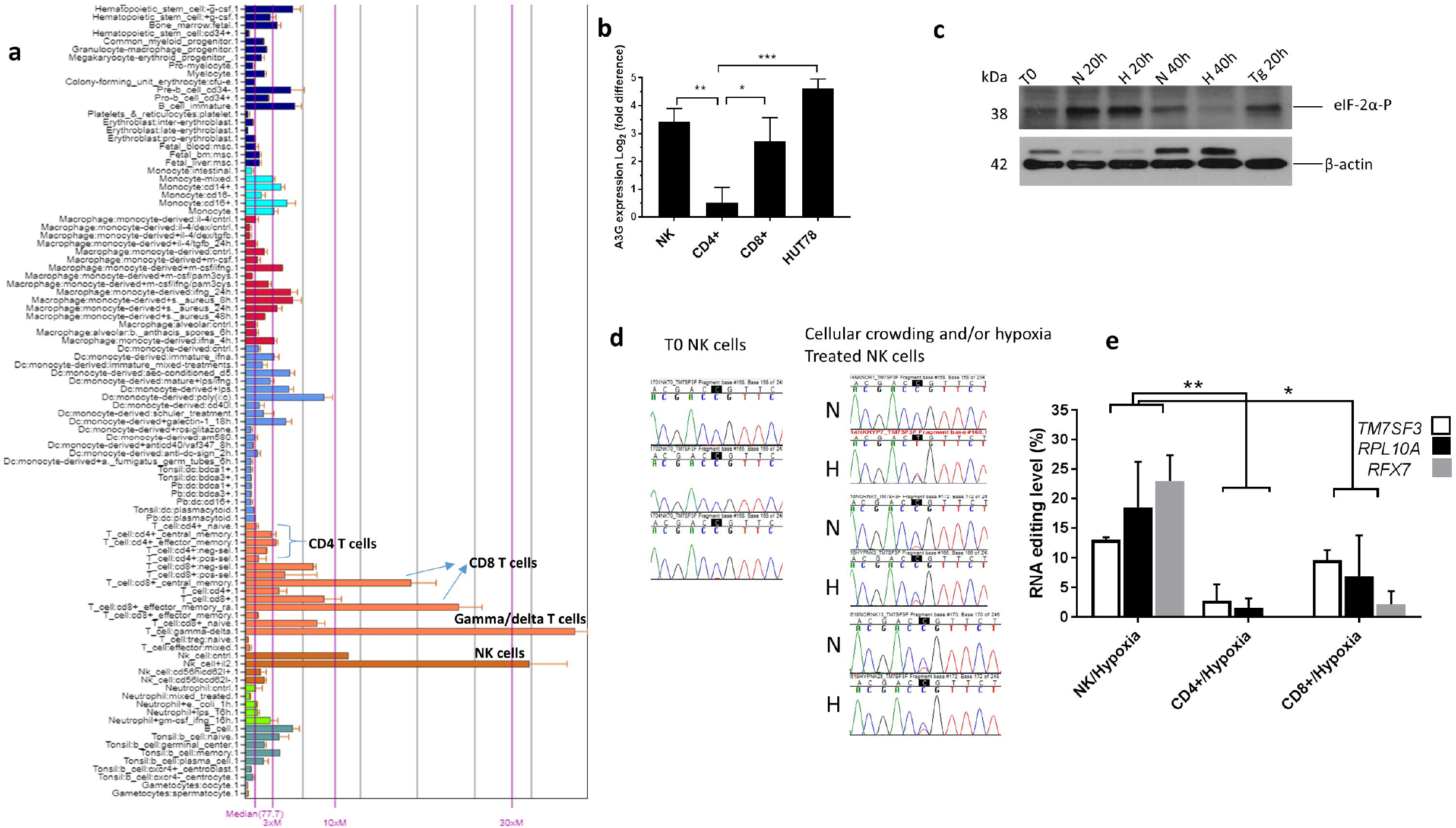
Cell specific expression of APOBEC3G (A3G) and the induction of RNA editing in NK cells. **(a)**Cell type specific expression of A3G (probe:214995_s_at) in Primary Cell Atlas, a metaanalysis of publicly available 100+ microarray datasets, available through the BIOGPS portal.**(b)***A3G* gene expression in NK, CD4+ T and CD8+ T cells. Gene expression measurements are normalized to that of β2-Microglobulin **(c)** Immunoblot showing the protein levels of eIF-2α phosphorylated at Ser 51 in whole cell lysates of NK cells at 0, 20 and 40 h under normoxia (N) or hypoxia (H). Thapsigargin (Tg) treated NK cells are used as a positive control and β-actin is used as a loading control **(d)** Sanger sequence chromatogram traces of cDNAs of PCR products of *TM7SF3* of unstressed (baseline,T0), normoxic (N) or hypoxic (H) NK cells. Edited C in *TM7SF3* is highlighted black **(e)** Estimation of site-specific C>U RNA editing by Sanger sequencing of RT-PCR products for *TM7SF3, RPL10A* and *RFX7* of NK, CD4+ T and CD8+ T cells subjected to hypoxia. See Methods for statistical analysis.

### RNASeq analysis of NK cells

To determine the transcriptome-wide targets of C>U RNA editing and their respective editing level in NK cells, we performed RNASeq analysis. PBMCs (n=3 donors) were cultured at a high density with/without hypoxia (1% O_2_) and site-specific editing of *TM7SF3* RNA was first confirmed, which showed higher level of editing in hypoxia relative to normoxia (Fig. 1d). The three normoxic and three hypoxic NK cells RNA samples were then sequenced by following the TruSeq RNA Exome protocol (see methods). Analysis of the RNASeq results was based on all C>U editing events in exons and UTRs that were (a) at least 5% in any sample, (b) overrepresented in the hypoxia group and (c) located in a putative RNA stem-loop structure [27] (see methods for details).

RNASeq analysis revealed 122 site-specific C>U editing events which were edited at a higher level in hypoxia as compared to normoxia, although editing also occurred in normoxia at variable levels due to cellular crowding in NK cells (Additional file 2; Table S1). The largest group of editing events comprised of non-synonymous changes, including 52 missense and 10 stop gain changes (Fig. 2a). Synonymous C>U editing events occurred in RNAs of 42 genes (Additional file 1; Figure S1b). We verified RNA editing by Sanger sequencing of cDNAs in 10 of 10 non-synonymously edited genes, which include *CHMP4B, EIF3I, FAM89B, GOLGA5, HSD17B10, RFX7, RPL10A, RPS2, TM7SF3* and *TUFM* (Fig. 2b). The highest level of non-synonymous RNA editing (~80%) occurred in *EIF3I*, which alters a highly conserved arginine to cysteine (c.C928T; R310C) (Fig. 2a and b). The average editing levels were lower for stop gain changes than for missense or synonymous changes and for changes in the UTRs and nc_RNA exonic sites, suggesting functional constraints on editing events that introduce stop-gain changes (Fig. 2c).

**Fig. 2.**
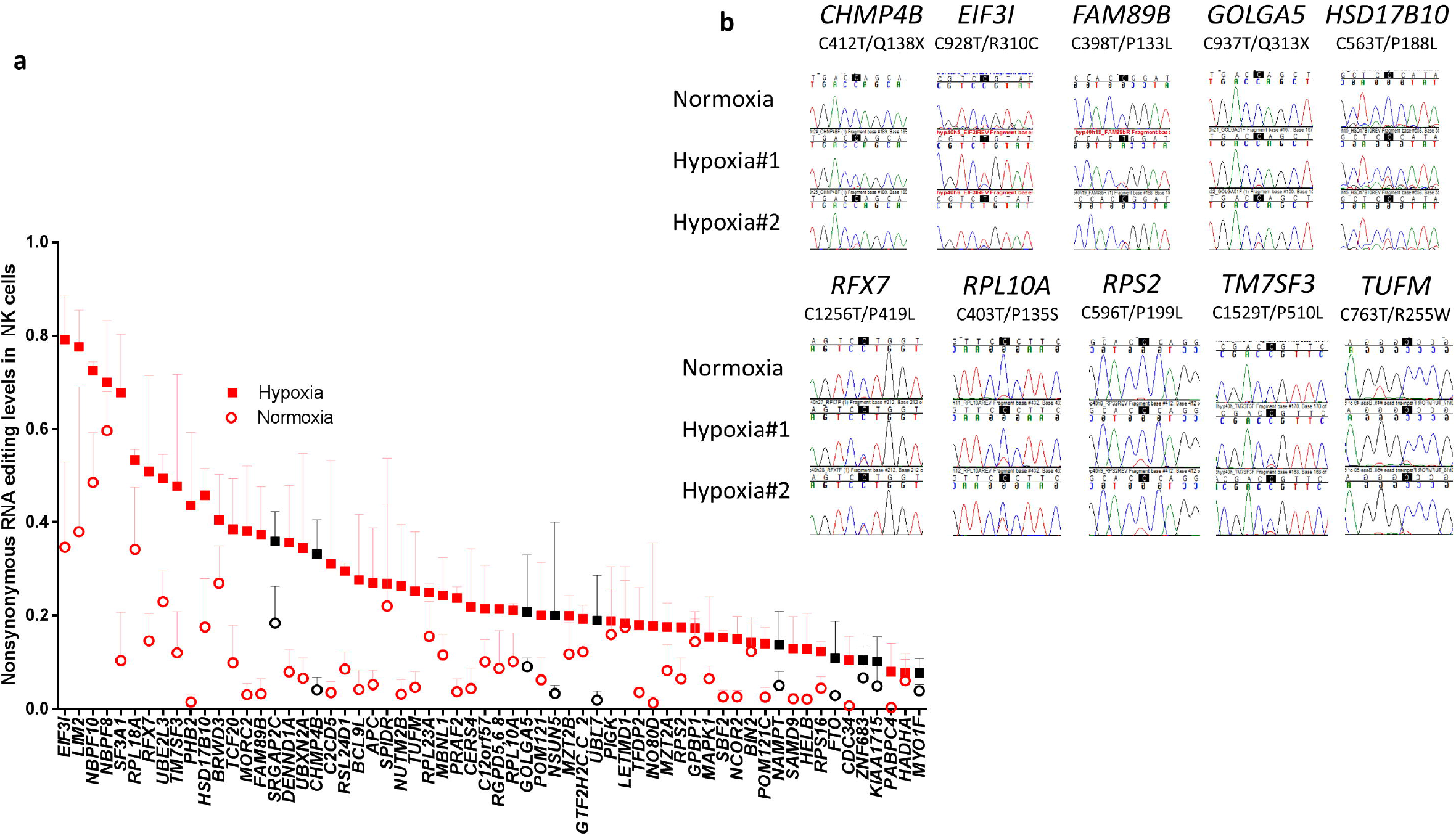
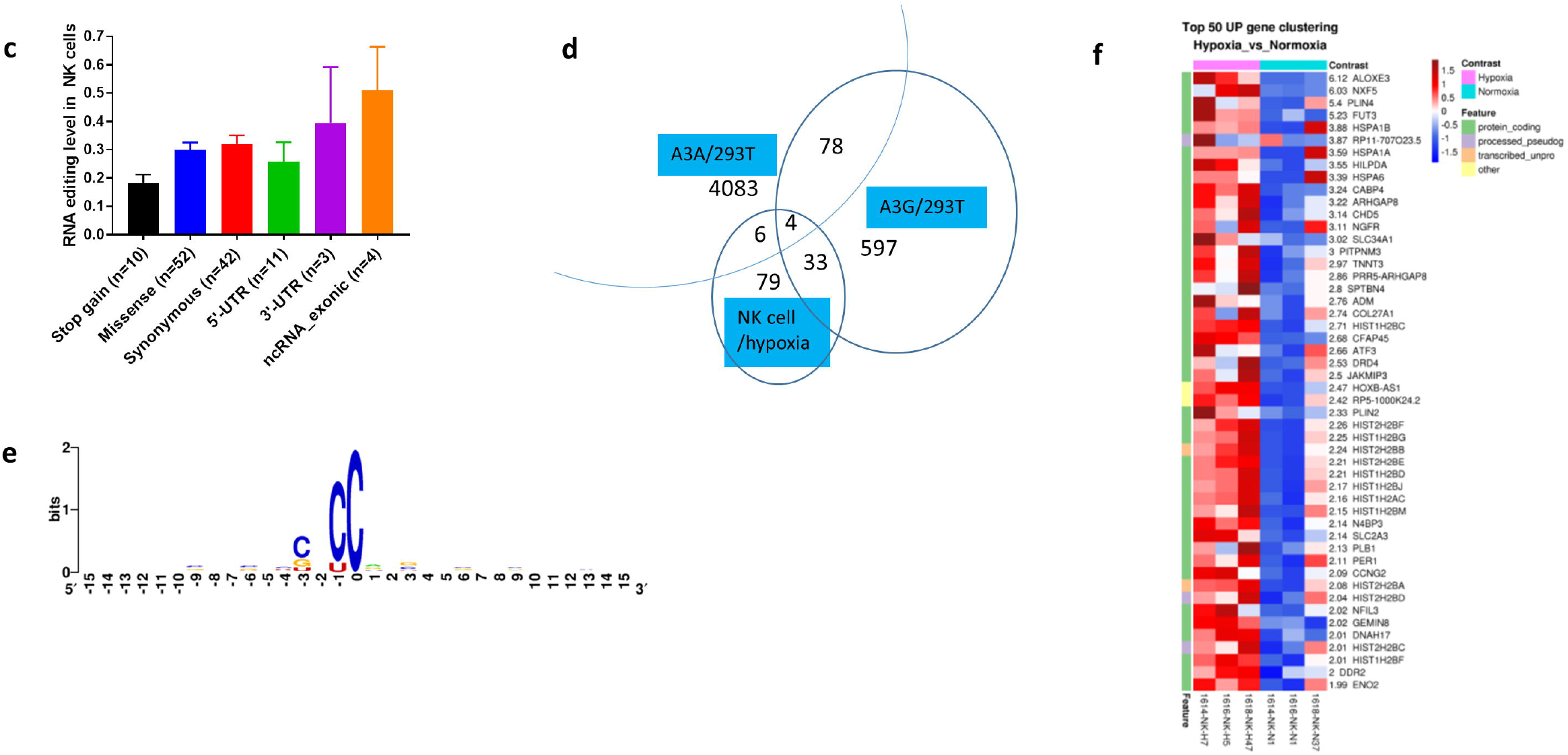
Distribution of site-specific A3G-mediated mRNA editing in NK cells. (a) A3G-mediated C>U RNA editing in NK cells resulting in non-synonymous changes (n=62) based in the order of highest to lowest editing level in hypoxia (40 h). Black symbols indicate genes that acquire nonsense RNA editing (n=10). (b) Sanger sequence chromatogram traces of amplified cDNA fragments comparing site-specific C>U editing in mRNAs of ten genes under normoxia and hypoxia. (c) Graph representing the editing levels of mRNA substrates of A3G in hypoxic NK cells and the location of editing in the mRNA as well as the type of change in the transcript sequence due to this editing (d) Venn diagram showing the number of unique and overlapping RNA editing sites (exonic and UTR) among hypoxic NK cells, 293T/A3A and 293T/A3G overexpression systems. (e) Logo indicating sequence conservation and nucleotide frequency for sequences bearing C>U editing sites (at position 0) among the edited transcripts in NK cells (n=122) (f) Heat map representing the most upregulated genes (n=50) in NK cells subjected to cellular crowding and hypoxia (cell stress)

We have previously identified the edited sites in RNAs of 293T/A3A and 293T/A3G overexpression systems [17, 28]. 37 of the 122 RNA editing sites in NK cells were among the 712 sites (exons+UTRs) in the 293T/A3G system (Additional file 3; Table S2) whereas only 10 edited sites in NK cells were among the 4,171 sites in the 293T/A3A system (Additional file 4; Table S3) (p=10^-5^, Fisher’s exact test) indicating that A3G is more likely to catalyze RNA editing in NK cells than A3A (Fig. 2d). Interestingly, 85 edited sites identified in NK cells did not overlap with those in the 293T/A3G system. Different parameters used for the computational analysis of edited sites, cell type specific factors and the method of induction of RNA editing (overexpression versus hypoxia) may play a role in the differences observed in the RNA editing targets of A3G in primary cells versus its exogenous overexpression in 293T cells. A3G has a preference for CC nucleotides both in its ssDNA and RNA substrates, whereas other A3 enzymes prefer TC nucleotides [13, 15, 17, 27, 28]. Sequence motif analysis of the 122 editing sites in NK cells shows a strong preference for C at -1 position (Fig. 2e), suggesting that these RNA editing events are catalyzed by A3G. The level of RNA editing and the expression of genes whose RNAs undergo editing show a weak positive correlation which is not statistically significant (r=0.1695, p=0.0620, n=122 genes, Additional file 1; Figure S2), suggesting that expression levels of the RNA edited genes do not influence RNA editing levels.

We determined the functional clustering of genes that undergo non-synonymous changes (n=62) due to RNA editing in NK cells using DAVID Bioinformatics Resources. The highest enrichment was for genes involved in “translation initiation”, “translation” and “ribosome” (Additional file 1; Figure S3) due to missense changes in RNAs of 8 genes (Table 1), including the highest non-synonymously edited *EIF3I* (Fig. 2a). RNA editing targeted highly conserved amino acids in 7 of 8 genes as predicted by at least 2 of the 3 softwares (Table 1; see Additional file 5; Table S4 for conservation analysis of all non-synonymous RNA editing sites). RNA editing also altered a conserved C (phyloP100 score=1.7811) at -4 nucleotide position in the 5’-UTR of another gene encoding the ribosomal protein, *RPLP0* (Additional file 2; Table S1). Since the regulation of translation plays a central role during cell stress [29, 30], these results suggest that RNA editing coordinately alters multiple ribosomal and other translational proteins, and may have an impact on the quality or quantity of protein translation under hypoxic stress.

**Table 1.**
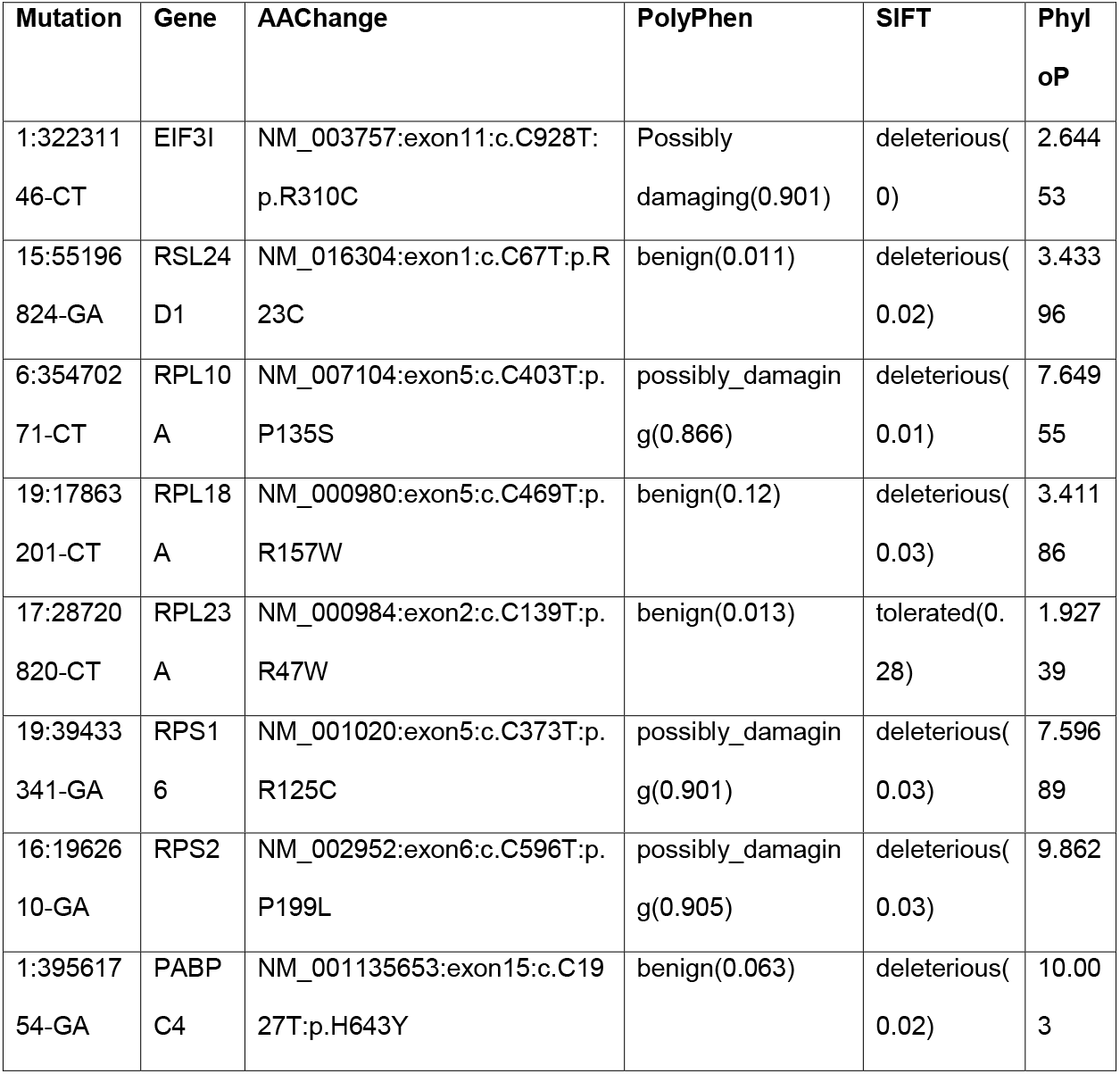
Conservation of amino acids recoded by A3G-mediated RNA editing in translational and ribosomal genesFigure Legends.

We also examined the changes in gene expression that occur during the induction of RNA editing in NK cells due to cellular crowding and hypoxia. We found upregulation of 82 genes and downregulation of 237 genes (fold change>2 and padj<0.005, Additional file 6; Table S5 and Additional file 1; Figure S4). Multiple genes of the heat shock protein HSP70 family (*HSPA1B, HSPA1A, HSPA6*) [31] and *ATF3*, which encodes a transcription factor integral to the ER stress response [32] are among the most upregulated (Fig. 2f). Thus, cellular crowding and hypoxia triggers a coordinated transcriptome remodeling in NK cells, which includes transcriptional induction of stress genes as well as recoding C>U RNA editing of translational and ribosomal genes.

### Confirmation of APOBEC3G-mediated RNA editing in lymphoma cell lines

To confirm A3G-mediated RNA editing and to examine the functional consequence of this editing in a cell line, we searched for cell lines that express A3G. We first examined the relative expression of A3G *in silico* in more than a 1,000 cell lines at the CCLE database. The highest expression of A3G was observed in leukemia and lymphoma cell lines (Additional file 1; Figure S5). Next, we ranked cell lines in the order of highest to lowest A3G expression (Fig. 3a) and selected JVM3 (rank=6), an EBV-transformed B cell prolymphocytic leukemia cell line, and HuT78 (rank=8), a CD4+ cutaneous T cell lymphoma cell line, which has also previously been used to identify A3G as a restriction factor in vif deficient HIV-1 viruses [16].

**Fig. 3.**
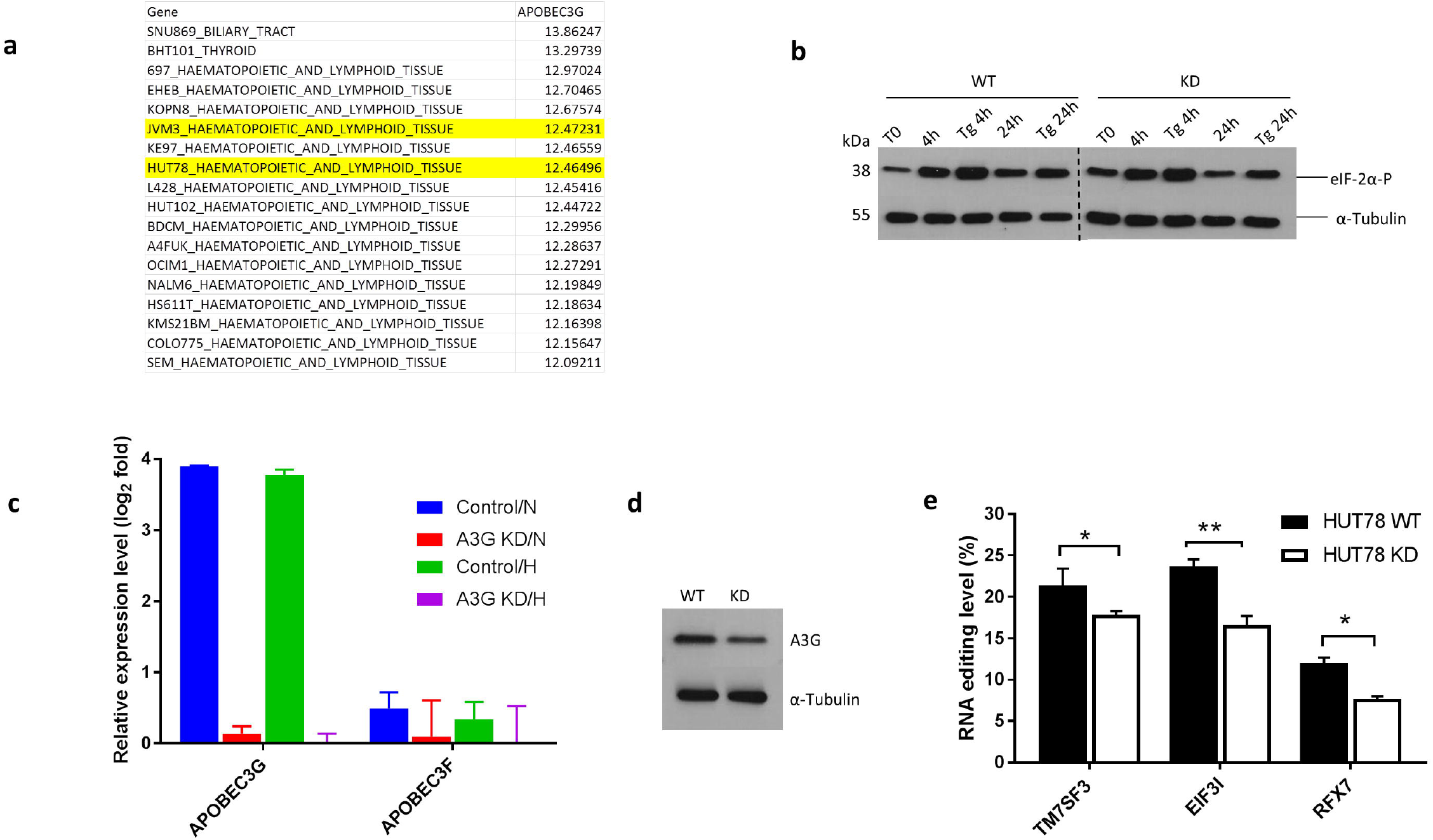
Distribution and induction of A3G-mediated C>U mRNA editing in lymphoma cell lines. **(a)** A List of cell lines in the CCLE database that have the highest expression of A3G (Affymetrix). The highlighted cell lines JVM3 and HuT78 are used in this study. **(b)** Immunoblot showing the protein levels of eIF-2α phosphorylated at Ser 51 in whole cell lysates of scramble WT and KD HuT78 cells at various time points. Thapsigargin (Tg) treated HuT78 cells is a positive control and α-Tubulin is used as a loading control. The WT and the KD HuT78 cells samples were run on two separate gels on the same day. The dashed line separates the two gels **(c)** *A3G* and *A3F* gene expression in control WT and KD HuT78 cells under normoxia (N) and hypoxia (H). Gene expression measurements are normalized to that of β2-Microglobulin (d) Immunoblot for A3G protein expression in whole cells lysates of WT and KD HuT78 cells. α-Tubulin is used as a loading control **(e)** Graph representing the percentage site-specific C>U RNA editing level for *TM7SF3, EIF3I* and *RFX7* of scramble WT and KD HuT78 cells in normoxia. See Methods for statistical analysis.

As compared to primary CD4+ T cells, A3G is highly expressed in the HuT78 lymphoma cell line (Fig. 1b and Fig. 3a). To further validate the RNA editing function of A3G, we knocked-down A3G in these cells using an A3G-specific shRNA lentiviral construct and a scramble negative control shRNA (referred to as WT HuT78). The WT HuT78 cells and the A3G knockdown cell line (KD) were further propagated and cultured at a high density of 1×10^6^ cells in 100 μl per well in a 96 well plate for 24 hours in normoxia or hypoxia (1% O_2_). High density culture and/or hypoxia treatment induced cell stress as there was an increased accumulation of phosphorylated eIF-2α [26], 4 h post culture (Fig. 3b). Under these conditions, we measured the expression of *A3G* in these cell lines by qPCR (Fig. 3c). The KD HuT78 cells showed markedly reduced expression of *A3G* as compared with WT HuT78 (Fig. 3c). We did not observe any significant variation in *A3G* levels with or without hypoxia treatment in the WT and KD HuT78 cells. *A3F*, which is also expressed in HuT78 did not show any significant variation in expression level between the WT and KD HuT78 cell lines, indicating that the knockdown for A3G is specific (Fig. 3c). We further confirmed the knock down of *A3G* by analyzing its protein expression by western blot (Fig. 3d) using specific antibodies to A3G. As compared to WT, KD HuT78 cells showed a reduction in A3G expression (Fig. 3d).

To determine the effect of A3G knock-down on RNA editing, we analyzed the editing level of three RNAs (*TM7SF3, EIF3I* and *RFX7*) previously validated as editing targets in NK cells. When cultured at a high density (mentioned above), we found site-specific editing of *TM7SF3, EIF3I* and *RFX7* RNAs in WT HuT78 cells and the level of editing was reduced in the A3G KD HuT78 cells (Fig. 3e and Additional file 1; Figure S6), correlating with the expression of A3G in these cells. We also confirmed the editing of *TM7SF3* in the JVM3 cell line in response to high cell density (Additional file 1; Figure S7a).

Considering that (1) A3G has a CC nucleotide preference, (2) RNA editing targets in NK cells and in 293T/A3G overexpression system overlap significantly (3) the same RNAs are site-specifically edited in NK and HuT78 cells-both highly expressing A3G; and (4) A3G KD HuT78 cells show decreased RNA editing, these results collectively indicate that A3G is an endogenous, inducible mRNA editing enzyme in NK, CD8+ and HuT78 (and JVM3) cells.

### A3G induces RNA editing by mitochondrial respiratory inhibition, independently of HIF-1α

To determine whether high density of HuT78 cells, which induces RNA editing by A3G, causes hypoxia, we cultured 1×10^6^ HuT78 cells in 100 μl per well in 96 well plates (high density) and the same number of cells in 1 ml culture in 6 well plates (low density), each under normoxia and hypoxia. We analyzed the stabilizatio of the hypoxia-inducible factor-1α (HIF-1α) protein, which is well known to be stabilized in hypoxic cells to promote the synthesis of mRNAs involved in cellular homeostasis [33], and measured the RNA editing levels of *TM7SF3*. As expected, HIF-1α was not stabilized at T0-when the cells were at a non-stressed state or under low density normoxic cell culture (6 well) after 24 hours (Fig. 4a). However, we found the stabilization of HIF-1α in cells cultured at a high density in 96 well plates both in normoxia and hypoxia, and in cells cultured at a low density in 6 well plates in hypoxia, suggesting that the high density 96 well normoxic culture had turned hypoxic (Fig. 4a). Under these conditions, RNA editing of *TM7SF3* was observed in cells cultured at a high cell density in both normoxia (20.6%) and hypoxia (20%) (Fig.4a; Additional file 1; Figure S7b). Although HIF-1α stabilization was observed in low cell density (6 well) hypoxic cultures, no RNA editing was observed under these conditions (Fig. 4a). These results confirm that as in NK cells RNA editing is induced by high cell density and hypoxia in HuT78 cells. Moreover, the stabilization of HIF-1α is not sufficient for the induction of RNA editing.

**Fig. 4.**
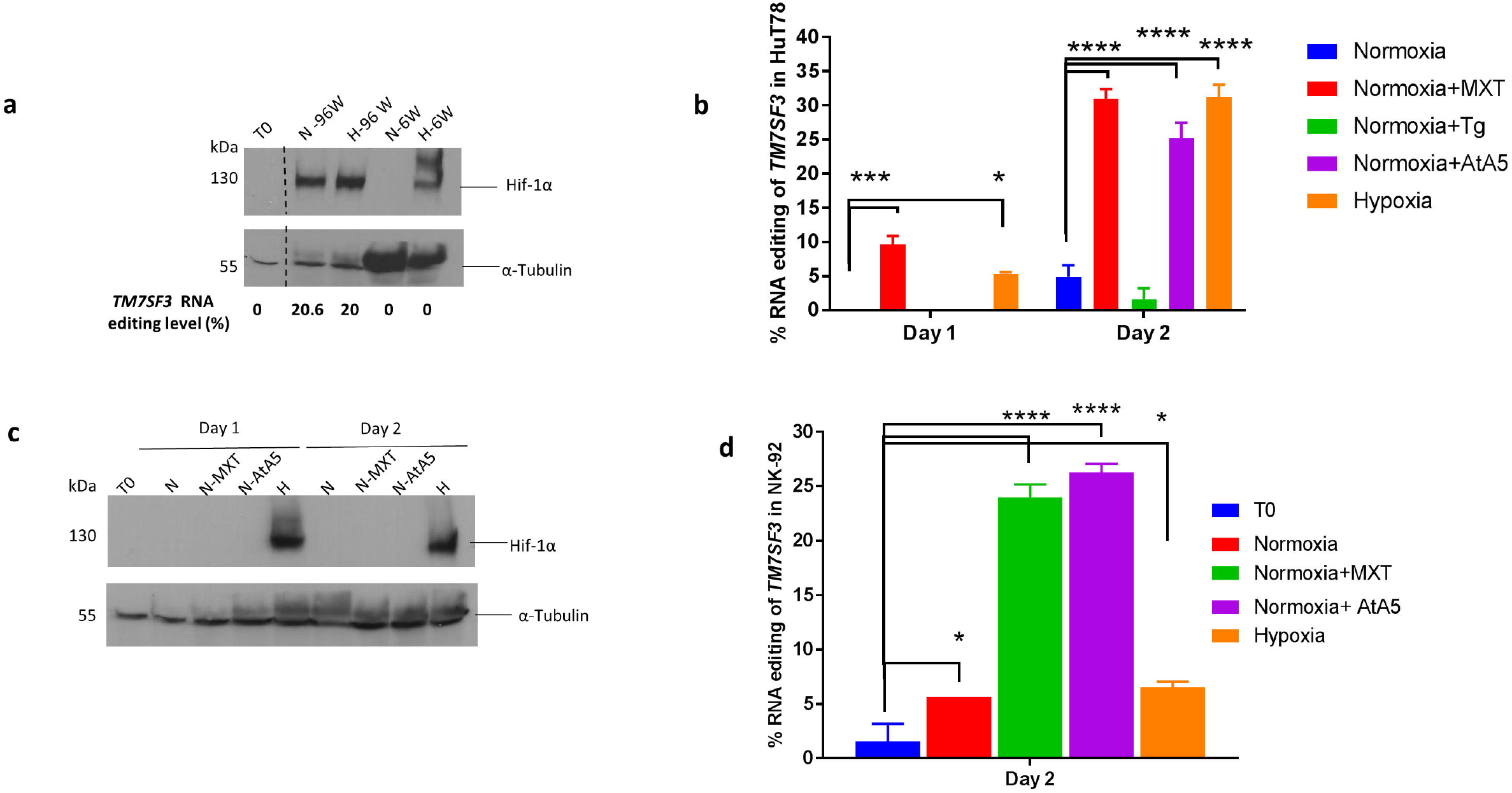
Induction of A3G-mediated C>U mRNA editing by the inhibition of mitochondrial respiration. **(a)**Immunoblot showing the protein level of HIF-1α in whole cell lysates of HuT78 when subjected to normoxia (N) and hypoxia (H) in 96 well (W) and 6 W plates for 24 hours. All lanes are part of the same gel. The dashed line represents the cropped region. The percentage C>U RNA editing levels in *TM7SF3* under these conditions is displayed below (b) The percentage C>U RNA editing in *TM7SF3* when HuT78 cells are treated with Myxothiazol (MXT), Thapsigargin (Tg), Atpenin (AtA5) and Hypoxia (H) for 24 hours (Day 1) or 42 hours (Day 2) (n=3) (c) Immunoblot showing the protein level of HIF-1α in whole cell lysates of HuT78 when subjected to normoxia (N) with or without the mitochondrial inhibitors (MXT and AtA5) and hypoxia (H) for one or two days (d) The percentage C>U RNA editing in *TM7SF3* when NK-92 cells are treated with Myxothiazol (MXT), Atpenin (AtA5) and Hypoxia (H) for 42 hours (n=3). See Methods for statistical analysis.

Previously we have shown that A3A-mediated RNA editing is induced by high cell density and hypoxia in hundreds of mRNAs in monocytes [15]. Furthermore, normoxic inhibition of the mitochondrial complex II by atpenin A5 (AtA5) and of the complex III by myxothiazol (MXT) mimic hypoxia and induce RNA editing as well as hypoxic gene expression in monocytes [34]. Since A3G-mediated RNA editing in NK and HuT78 cells is also induced by hypoxia, we tested the effect of these mitochondrial inhibitors on RNA editing in HuT78 cells cultured in normoxia. Additionally, to test whether endoplasmic reticulum (ER) stress can also induce RNA editing, we treated the cells with Thapsigargin (Tg). Tg induces ER stress by raising intracellular calcium levels and lowers the ER calcium levels by specifically inhibiting the endoplasmic reticulum Ca^++^ ATPase [35, 36], resulting in the accumulation of unfolded proteins and an increased accumulation eIF-2α phosphorylated at Ser 51 (Figs. 1c and 3b). To test the effect of hypoxic stress alone on HuT78 cells, we reduced the cell density to avoid cellular crowding and cultured the cells at an intermediate density of 0.5×10^6^ cells per 500 μl per well in 24 well plates with or without the chemical inhibitors in normoxia, and hypoxia alone for one or two days. Under these conditions, we determined RNA editing level and the stabilization of HIF-1α in these cells. We observed that RNA editing is mildly induced in cells treated with MXT and by hypoxia alone on day 1, at approximately 10% and 5% levels, respectively (Fig. 4b). RNA editing levels increased to approximately 30% in cells treated with MXT, AtA5 or hypoxia alone on day 2. Treatment of cells with Tg did not induce RNA editing (Fig. 4b). Furthermore, HIF-1α was stabilized only when the cells were subjected to hypoxia but not in normoxia in the presence or absence of the mitochondrial inhibitors (Fig. 4c). These results suggest that RNA editing induced by hypoxic stress at a high cell density is triggered by mitochondrial respiratory inhibition and occurs independently of the stabilization of HIF-1α as well as the ER stress response.

Although the A3G expression data did not include the NK-92 lymphoma cell line in the CCLE database, given its similar characteristics to primary NK cells and the convenience of culturing NK-92 cells as compared with primary NK cells, we tested the induction of RNA editing in NK-92 cells. We treated NK-92 cells with normoxia with or without the mitochondrial inhibitors (AtA5 or MXT) or hypoxia alone at intermediate density in 24 well plates for 2 days. Interestingly, RNA editing was induced by the inhibition of mitochondrial respiration (~25%), but only slightly by hypoxia treatment (Fig. 4d) in NK-92 cells. The reason behind the difference in hypoxia induced RNA editing level of HuT78 and NK-92 cells may be due to metabolic differences between the two cell lines. However, the induction of A3G-mediated RNA editing due to mitochondrial respiratory stress in NK-92 cells provides a model system and an opportunity for further functional studies.

### APOBEC3G promotes Warburg-like metabolic remodeling and suppresses proliferation under stress

We have previously identified *SDHB* and *SDHA* mitochondrial complex II subunits as targets of A3A-mediated RNA editing in hypoxic monocytes [15]. In the current study, we find that A3G non-synonymously edits several mitochondrial genes’ RNAs including *TUFM, HADHA, HSD17B10* and *PHB2* in hypoxic NK cells (Fig. 2a). Thus we hypothesized that hypoxic stress-induced RNA editing by A3G alters mitochondrial function.

To test the role of A3G on bioenergetics in response to high cell density and hypoxic stress, we measured the metabolic profile of WT and KD HuT78 cells using the Seahorse platform. We performed the mitochondrial and the glycolytic stress tests to measure the oxygen consumption rate, representative of basal respiration and the extracellular acidification rate, representative of glycolysis in cells cultured at a high density in three separate experiments (Fig. 5a). We have presented metabolic alterations as respiration-to-glycolysis ratio (R/G) both in unstressed (T0) and stressed cells (Fig. 5b). As expected, cell stress caused by high cell density reduced R/G ratio in each experiment relative to unstressed T0 cells, indicating a decrease in respiration relative to glycolysis. However, R/G ratios decreased to a lesser extent under stress in A3G KD, relative to WT HuT78 cells, indicating that A3G plays a role in reducing mitochondrial respiration relative to glycolysis under hypoxic stress caused by high cell density (Fig. 5b).

**Fig. 5.**
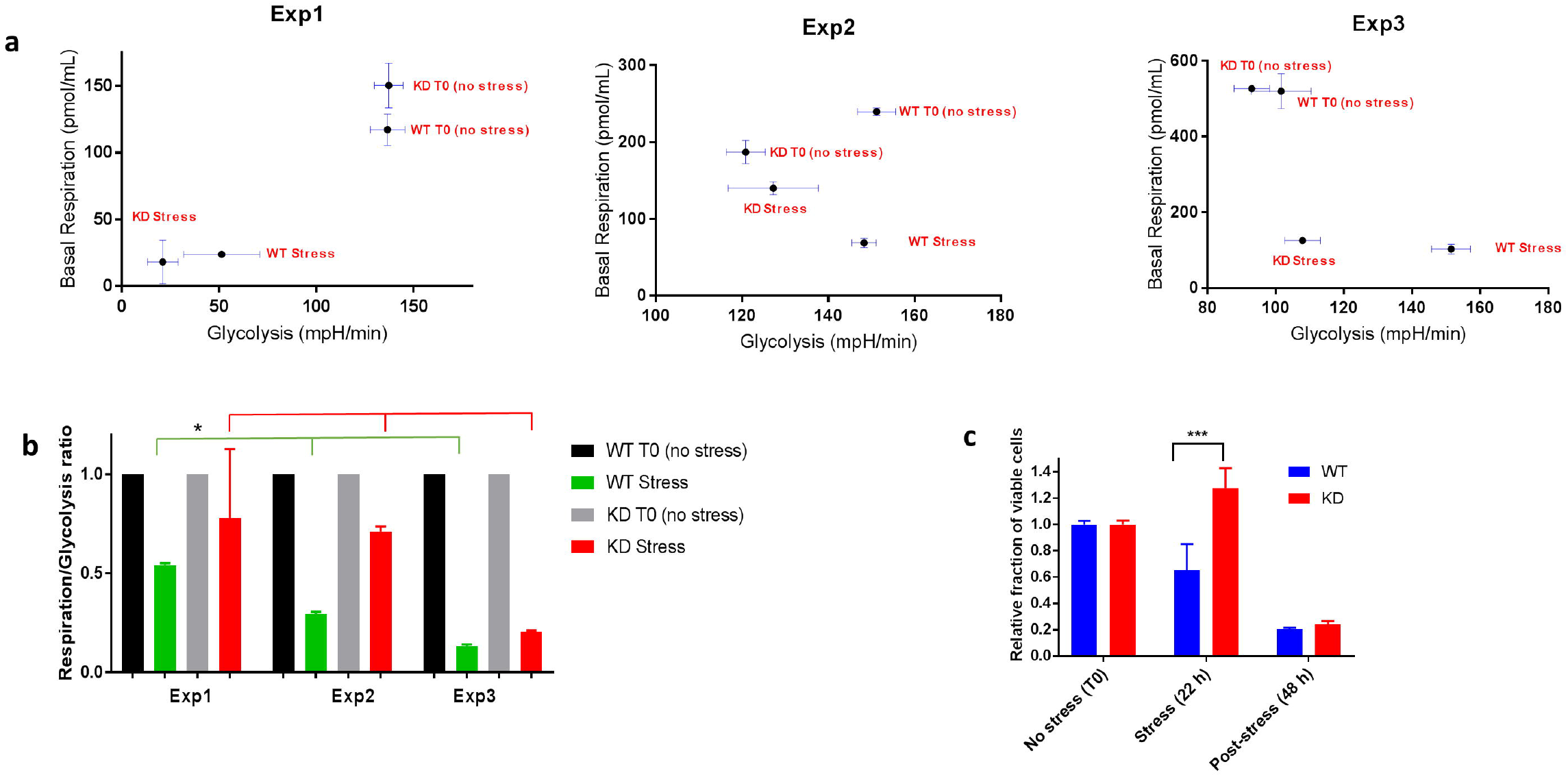
A3G-mediated C>U mRNA editing results in Warburg-like effect in lymphoma cell lines. (a) Plot representing the basal respiration versus glycolysis in T0 unstressed and stressed cells (cellular crowding in normoxia) in WT and KD HuT78 cells (mean and SD, n=3-4). (b) Bar graph showing the respiration to glycolysis ratios (R/G) normalized to unstressed WT and KD HuT78 cells are shown (mean and SEM). (c) Bar graph representing the fraction of viable WT and KD HuT78 cells when subjected to cellular crowding for 24 hours in normoxia (see methods) followed by culture in non-stressed conditions for another 48 hours (Mean and SEM, n=3). See Methods for statistical analysis.

Hypoxic stress can suppress translation and lead to growth arrest by inhibiting cell cycle progression in non-transformed cells or by promoting apoptosis by the p53 pathway in transformed cells [29, 37]. To examine the role of A3G on cellular proliferation under stress, we measured the proliferation of the WT and KD HuT78 cells when cultured at a high density for 22 hours followed by ‘recovery period’ by culturing these stressed cells at a low density for another 48 hours. The fraction of viable cells reduced in WT, but increased in A3G-KD HuT78 cells during 22 hours of stress (Fig. 5c) (mean±SEM=0.653±0.197 versus 1.277± 0.151; n=3), indicating that A3G-KD HuT78 cells proliferated more under high density culture conditions. However, the number of viable cells in WT and A3G-KD HuT78 cells did not show any difference at 48 hours after recovery from stress when cultured under non-stress conditions (Fig. 5c). These results suggest that hypoxic stress in lymphoma cells suppresses proliferation in vitro and that A3G plays an important role in this suppression.

## Discussion

In this study we find that A3G edits scores of RNAs in NK cells and CD8+ T lymphocytes as well as lymphoma cell lines, when cultured at a high density and hypoxia. A3G-mediated site-specific RNA editing is triggered by the inhibition of mitochondrial respiration, and targets the mRNAs of many ribosomal and translational genes resulting in non-synonymous changes. A3G reduces mitochondrial respiration relative to glycolysis, and suppresses cell proliferation under stress in transformed lymphoma cells (Fig. 6). These results identify A3G cytidine deaminase as the third endogenous C>U RNA editing enzyme in mammals and together with A3A in myeloid cells, defines a new functional category of RNA editing enzymes that are active in immune cells. In addition, our findings uncover a previously unrecognized gene regulation mechanism in NK and CD8+ T cells that is induced by hypoxic stress.

**Fig. 6.**
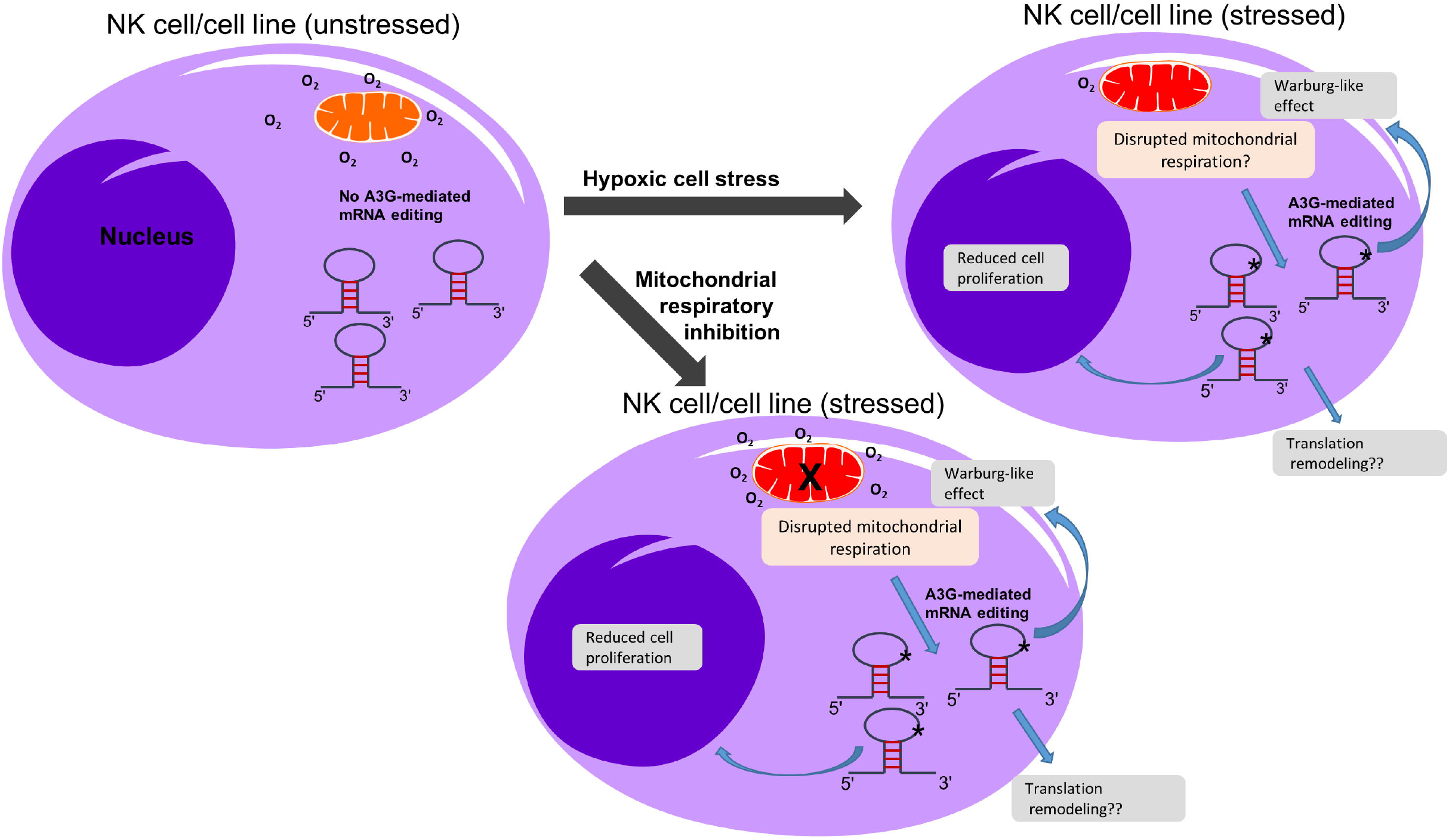
Simplified diagram summarizing the induction and relevance of A3G-mediated site-specific C>U cellular mRNA editing in NK cells and lymphoma cell lines. NK / lymphoma cell is shown under normal physiological conditions when the cells are unstressed (left) or when the cells are stressed by hypoxia (top right) or due to the inhibition of mitochondrial respiration (bottom right). Under normal physiological conditions (baseline) mRNAs (stem-loop) in NK cells do not undergo C>U RNA editing. Under hypoxic stress or upon mitochondrial respiratory inhibition, an unknown signal originating in the mitochondria (red) triggers site-specific A3G-mediated C>U editing in multiple mRNA substrates bearing a stem-loop structure. The cellular mRNA editing induced by mitochondrial hypoxic stress may result in translational reprogramming of NK cells, Warburg-like metabolic remodeling by preferring glycolysis over mitochondrial respiration, and reduced cellular proliferation in order to promote adaption during NK/lymphoma cell stress.

There are two major differences in A3-mediated RNA editing and ADAR- and APOBEC1-mediated editing. First, A3-mediated RNA editing is induced upon hypoxic stress (A3A and A3G) or by IFNs (A3A), while it is essentially absent or rare in baseline unstressed immune cells[15](Fig. 1d). In contrast, ADAR and APOBEC1-mediated RNA editing events occur in baseline unstimulated cells [38-40]. Second, A3-mediated RNA editing events occur in exonic coding regions of genes as commonly as in UTRs [15] (Fig. 2c), whereas ADAR- and APOBEC1-mediated RNA editing events preferentially occur in UTRs, where they are at least an order of magnitude more frequent relative to coding exons [38-40]. Together, these findings suggest that A3-mediated RNA editing plays a role in response to certain cell stress by altering protein function.

A recurrent theme in many types of cell stress responses, including ER and mitochondrial unfolded protein stress response generally caused by heat shock, nutrient deprivation, hypoxia or DNA damage, is the regulation of gene expression. This is achieved by the general suppression or reprogramming of translation to promote recovery from stress or cell death [30, 41]. The highest level of RNA editing resulting in a non-synonymous change is observed in *EIF3I* in hypoxic NK cells. *EIF3I* encodes a subunit of EIF3, the most complex translation initiation factor comprised of 13 subunits in mammals, which is involved in all molecular aspects of translation initiation. The EIF3 complex has been implicated in the translation of mRNAs important for cell growth [42] and mitochondrial respiration [43], and its subunits are overexpressed in multiple cancers [44]. Interestingly, EIF3I was previously shown to have decreased protein synthesis in cold-stressed mammalian cells, implying its important role in stress response and recovery [45]. Consistent with these reports, we find that the knockdown of A3G in HuT78 lymphoma cells reduces the predicted deleterious RNA editing of *EIF3I* in association with reduced mitochondrial respiration and cell proliferation (Additional file 5; Table S4) during hypoxic stress. Thus, our findings suggest that A3G promotes hypoxic stress responses via RNA editing of *EIF3I*, ribosomal/translational genes and possibly other stress-related genes.

Cancer cells switch to aerobic glycolysis even in the presence of a functional mitochondria and this phenomenon is termed the ‘Warburg effect’. However, the function of Warburg effect in tumor growth, proliferation and support of cellular biosynthetic programs is still inconclusive [46]. In response to acute hypoxia, A3G-medited RNA editing in the WT cells may promote Warburg effect by preferring glycolysis over mitochondrial respiration and decreased translation, while limiting overall cellular proliferation.

Interestingly, even though normal B cells and plasma cells show low expression of A3G (Fig 1a), we find that the highest expression levels are observed in neoplastic B and plasma cell lines derived from acute lymphoblastic leukemia, B-cell lymphoma, Burkitt lymphoma and multiple myeloma. Increased expression of A3G in many B-cell leukemia/lymphoma cell lines (Fig. 3a), and NK/T cell lymphoma [47] supports the notion that it may play an oncogenic role by enhancing survival under oxygen-limiting conditions. It is known that NK cell function is impaired in the tumor micro-environment or chronic infections due to multiple factors, including hypoxia [48]. This may be achieved in part by A3G-mediated RNA editing resulting in the cellular remodeling during stress.

We also find that RNA editing by A3G can be induced by normoxic inhibition of mitochondrial respiration and occurs independently of HIF-1α stabilization (Fig. 4), in a manner similar to the regulation of A3A-mediated RNA editing in monocytes [34]. Earlier studies have shown that the inhibition of mitochondrial respiration antagonizes the stabilization of HIF-1α in hypoxia [49]. Despite the lack of HIF-1α stabilization, however, we find that mitochondrial respiratory inhibition mimics hypoxia and induces RNA editing by A3G. Thus, A3G-mediated RNA editing joins a growing number of hypoxia-induced responses that can be mimicked by the inhibition of mitochondrial respiration. These include carotid body paragangliomas caused by mitochondrial complex II mutations [50], expression of hypoxia-related genes and A3A-meditated RNA editing responses in monocytes [34], stimulation of the cardiorespiratory system by carotid body chemoreceptors [51], hypoxic pulmonary vasoconstriction mediated by pulmonary arterial smooth muscle cells [52], and hypoxia-induced changes in astrocytes [53], the most abundant glial cells in the brain. We hypothesize that hypoxia triggers A3G-mediated RNA editing downstream of a pathway activated by mitochondrial respiratory inhibition as a result of severe oxygen deprivation or respiratory inhibitors in normoxia (Fig. 6). Details of this mitochondrial hypoxic signaling pathway that activates A3-mediated RNA editing are subject of future studies.

Finally, the unexpected discovery of RNA editing functions for A3A and A3G require reconsideration of the physiological functions of the A3 enzymes solely as anti-viral factors. For example, A3G evolved with positive selection signature for millions of years in the primate lineage before humans were infected by HIV-1 [54]. Also, A3G orthologs that have the signature of positive evolutionary selection are present in primates that are not infected by SIVs [55]. Although suppression of endogenous retroviruses was speculated as an *in vivo* function of A3 enzymes, mouse A3 knockout is viable without any evidence of catastrophic retroviral infection [56]. Furthermore, the anti-HIV model of the double-domain A3G does not adequately explain why the zinc-coordinating residues in the N-terminal domain are conserved, since ssDNA deamination of HIV-1 minus strand by A3G in target cells does not require catalytic activity of the N-terminal domain [13, 14]. In contrast, RNA editing requires the conserved zinc-coordinating residues in both its N-and C-terminal domains [17]. Thus, cellular RNA editing provides a plausible explanation for A3G’s long-term evolutionary history, the presence of two conserved zinc-coordinating catalytic domains and the high expression patterns in NK and CD8+ T cells. In conclusion, our findings suggest that the primary function of A3G *in vivo* may be cellular RNA editing to facilitate adaptation to mitochondrial hypoxic stress in lymphocytes. Further studies are required to examine the RNA editing function of the other APOBEC3 enzymes, as well as their significance in immunity.

## Conclusion

This study shows the endogenous inducible site-specific RNA editing activity of the A3G cytidine deaminase, the most studied member of the APOBEC3 family, and suggests its physiological function in human immune and transformed cells. Widespread RNA editing by A3G can facilitate cellular adaptation to hypoxic cell stress triggered by mitochondrial respiratory inhibition in primary cytotoxic lymphocytes and lymphoma cell lines. A3G is the third endogenous C>U RNA editing enzyme to be identified in mammals. In addition, our study uncovers a novel epitranscriptomic gene regulation mechanism in cytotoxic lymphocytes, specifically NK cells. APOBEC3 cytidine deaminases may define a new class of RNA editing enzymes that are induced in response to certain cell stress factors.

## Methods

### RNA Sequencing

RNAs (DNA-free) were extracted from NK cells of 3 donors subjected to normoxia and hypoxia treatments (6 samples total) using the Total RNA clean-up and concentration kit (Norgen Biotek) as per the manufacturer’s instructions. RNA Libraries were prepared using the Illumina TruSeq RNA Exome protocol and kit reagents. RNA input for intact total RNA was 10 ng. RNA QC analysis by electrophoresis (2100 Expert, B.02.08.SI648, Agilent Technologies, Inc.) showed RIN numbers of 9.6, 7.8, 6.4 for normoxic and 2.8, 9.4 and 2 for hypoxic samples. These RIN numbers showed evidence of RNA degradation. Therefore, for degraded RNA samples input amount was determined by calculating the percentage of RNA fragments >200 nt (DV200) by running the samples on an RNA ScreenTape (Agilent Technologies) and performing region analysis using the Tapestation Analysis Software. Based on the DV200 calculation of 52-85%, 40 ng was the input amount and was considered suitable for this protocol. Fragmentation of the RNA was performed on intact samples. First and second strand synthesis were preformed to generate double-stranded cDNA. The 3’ ends were adenylated and Illumina adapters were ligated using T-A ligation. PCR was performed to generate enough material for hybridization and capture. PCR products were validated for the correct sizing using D1000 Screentape (Agilent Technologies). 200 ng of each product was pooled together in 4-plex reactions for hybridization and capture. Two sequential rounds of hybridization and capture were performed using the desired Capture Oligo pool. A second round of PCR was done to generate sufficient libraries for sequencing. Final libraries were validated for correct size distribution on a D1000 Screentape, quantified using KAPA Biosystems qPCR kit, and the 4-plex capture pools were pooled together in an equimolar fashion, following experimental design criteria.

Each pool was denatured and diluted to 2.4 pM with 1% PhiX control library added. Each pool was denatured and diluted to 16 pM for On-Board Cluster Generation and sequencing on a HiSeq2500 sequencer using 100 cycle paired-end cluster kit and rapid mode SBS reagents following the manufacturer’s recommended protocol (Illumina Inc.) and 100 million paired reads per sample were obtained.

### RNA editing bioinformatics analysis

*RNA editing events detection:* Sequence reads passing quality filter from Illumina RTA were first checked using FastQC [57] and then mapped to GENCODE(https://www.gencodegenes.org/) annotation database (V25) and human reference genome (GRCh38.p7) using Tophat2 [58] with a lenient alignment strategy allowing at most 2 mismatches per read to accommodate potential editing events. The mapped bam files were further QCed using RSeqQC [59]. Then all samples were run through the GATK best practices pipeline of SNV calling (https://gatkforums.broadinstitute.org/gatk/discussion/3892/the-gatk-best-practices-for-variant-calling-on-rnaseq-in-full-detail) using RNASeq data to obtain a list of candidate variant sites. All known SNPs from dbSNP (V144) [60] were removed from further analyses.

### Hypoxia induced editing events filtering

Pileups at candidate sites were generated using samtools for all samples and the base counts for alternative and reference base were calculated. Potential candidates for RNA editing were first filtered using the following two criteria: (a) at least 5% editing level on any sample within the population; (b) only C>T and G >A events were selected. The editing base counts were modeled as Binomial distribution and the effect of hypoxia on RNA editing at each site was tested with a generalized linear model (GLM) using paired samples. Multiple test adjustment was applied using Benjamini-Hochberg procedure to control false discovery rate (FDR). Hypoxia induced editing events were identified with log-odds-ratio greater than 0 and adjusted-p value less than 0.05.

### Results

A table specifying the editing site, the type of editing event, editing level and number of reads on a reference and alternative bases on each sample for each group was initially produced filtering events with OR > 1 and a FDR < 0.05 level.

### Annotation

Hypoxia induced editing events passing filters were annotated using ANNOVAR [61] with RefSeq gene annotation database to identify gene features, protein changes and potential impact. Also 15 base pair upstream and downstream flanks from the variant sites were displayed in separate columns.

### Manual filter

The above analyses initially revealed 383 C>U editing sites which were then subjected to a final stringent manual filtering step which retained only those sites (a) in exons and UTRs, (b) with-1 position (relative to edited C) is either a C or T and (c) within a stem-loop structure where the edited C is at the 3’-end of a putative tri- or tetra loop which is flanked by a stem that was at least 2 base pair long when base complementarity was perfect, or at least 4 base pair long when complementarity was imperfect by 1 nucleotide mismatch or 1 nucleotide bulging. This stringent manual filter reduced the number of edited sites to 122 (Additional file 2; Table S1).

### RNASeq differential expression analysis

Raw counts for each gene were generated using HTSeq [62] with intersection_strict mode. Differential gene expression was analyzed by DESeq2 [63]. Bioconductor package with paired sample design to identify hypoxia induced gene expression changes.

### Conservation analysis of amino acids recoded by RNA editing in NK cells

The impact of non-synonymous RNA editing on protein function was examined by PolyPhen and SIFT programs from ENSEMBL VEP tool, which give a score and a verbal description of the impact (https://useast.ensembl.org/info/docs/tools/vep/index.html). In addition, conservation score based on 100 vertebrates basewise conservation was obtained from UCSC (phyloP100way).

### Isolation and culture of cells

The HuT78,JVM3 and NK-92 cell-lines were obtained from ATCC. Hut 78 cells were cultured in IMDM (ATCC) containing 20% Fetal Bovine serum (FBS) (Sigma-Aldrich), JVM3 cells were cultured in RPMI (ATCC) containing 10% FBS and NK-92 cells were cultured in Alpha Minimum Essential medium without ribonucleosides and deoxyribonucleosides (Life Technologies) but with 2 mM L-glutamine and 1.5 g/L sodium bicarbonate as well as 0.2 mM inositol, 0.1 mM 2-mercaptoethanol, 0.02 mM folic acid, 500 U/ml IL-2 (Aldesleukin - a kind gift from Novartis), 12.5% horse serum (ATCC) and 12.5% FBS. Peripheral blood mononuclear cells (PBMCs) of anonymous platelet donors were isolated from peripheral blood in Trima Accel™ leukoreduction system chambers (Terumo BCT) in accordance with an institutional review board-approved protocol, as described earlier [15], in RPMI-1640 medium (Mediatech) with 10% FBS, 100 U/ml penicillin and 100 μg/ml streptomycin (Mediatech). NK, CD4+ and CD8+ cells were isolated from PBMCs (cultured at 5×10^7^ in 1.8 ml per well in 12 well plates) by immunomagnetic negative selection using the EasySep™ Human NK Cell Isolation Kit (Stemcell Technologies, catalog # 17955), EasySep™ Human CD4+ Cell Isolation Kit (Stemcell Technologies, catalog # 17952) and EasySep™ Human CD8+ Cell Isolation Kit (Stemcell Technologies, catalog # 17953), respectively, following the manufacturer’s instructions. Enrichment for NK cells was > 90% (Additional file 1; Figure S8) and that of CD4+ and CD8+ was >99%, as verified by flow cytometry.

### Cell stress and inhibitor treatment

For cell crowding experiments, the HuT78 cells were cultured at a density of 0.5-1×10^6^ cells per 100 μl per well in 96 well plates for 22-24 hours at 37° C.

For hypoxia treatment, PBMCs were cultured at a density of 5×10^7^ in 1.8 ml per well in 12 well plates under 1% O_2_, 5% CO_2_ and 94% N_2_ in an Xvivo™ System (Biospherix) for 40 hours. Following culture, NK, CD4+ and CD8+ cells were separated as mentioned above. In case of HuT78, the cells were cultured in the hypoxia chamber for 24 or 40 hours at a density of 1×10^6^ cells per ml in 6 well plates.

For testing the mitochondrial inhibitors, HuT78 and NK-92 cells were cultured at 0.5×10^6^ cells per 0.5 ml in 24 well plates in normoxia with or without AtA5 and MXT or hypoxia alone for 2 days at 37°C.

Human IFN-γ was obtained from PeproTech and used at a concentration of 50 ng/ml. AtA5 (Cayman chemical #11898) and MXT (Sigma Aldrich #T5580) was used at a concentration of 1 μM.

### Extracellular Flux Assays

HuT78 cells (scramble WT and KD) were plated in 96-well plates at a density of 0.5 or 1×10^6^ in 100 μl per well (total 3×10^6^ cells) and incubated for 22-24 hours at 37⁰C. The cells were harvested and washed with PBS and re-counted on a hemocytometer (INCYTO C-Chip). Half of the cells were re-suspended in the XF base media specific for the Mitochondrial and the other half in XF base media specific for the Glycolytic Stress Tests (below), respectively. For all extracellular flux assays, cells were plated on cell-tak coated Seahorse XF96 cell culture microplates in (duplicate, triplicate or quadruplicate, depending on the cell count post culture) at a density of 3-6 X10^5^ cells per well. The assay plates were spin seeded for 5 minutes at 1,000 rpm and incubated at 37°C without CO_2_ prior to performing the assay on the Seahorse Bioscience XFe96 (Agilent). The Mitochondrial Stress Test was performed in XF Base Media containing 10 mM glucose, 1 mM sodium pyruvate, and 2 mM L-glutamine and the following inhibitors were added at the final concentrations: Oligomycin (2 μM), Carbonyl cyanide 4-(trifluoromethoxy)phenylhydrazone (FCCP) (2 μM), Rotenone/Antimycin A (0.5 μM each). The Glycolytic Stress Test was performed in XF Base Media containing 2 mM L-glutamine and the following reagents were added at the final concentrations: Glucose (10 mM), Oligomycin (2 μM), and 2-deoxy-glycose (50 mM).

### shRNA-mediated Knock-down of APOBEC3G in HuT78 cells

A3G knock-down in Hut78 cells was performed at the RPCCC gene modulation shared resource. For A3G knock-down, GIPZ human A3G shRNAs with the following Clone ID’s were used: V2LHS_80856, V2LHS_80785, V2LHS_80786 (Dharmacon). Lentiviruses were produced by cotransfection of 293T cells with A3G shRNA (or pGIPZ non-silencing control) along with psPAX2 and pMD2.G packaging plasmids, using the LipoD293 reagent (1:2.5 DNA to lipoD293 ratio) (SignaGen Laboratories) as per the manufacturer’s instructions. Culture supernatants were collected 48 and 72 hours after transfection and cleared by filtration through 0.45 μm cellulose acetate syringe filter. For shRNA expression, 1×10^6^ Hut78 cells were pelleted and resuspended with 1 ml culture supernatants containing the virus and 1 μl of 4 mg/ml polybrene. The cells were placed in 6 well plates and incubated for 30 mins at 37° C. The plate was sealed and spun at 1800 rpm for 45 mins in a microtiter rotor (Beckman Coulter) at room temperature and then incubated for 6 hours at 37 °C. After infection the cells were centrifuged at 500g for 5 mins and resuspended in IMDM media and incubated for 48 hours at 37° C. Puromycin (1 μg/ml) was added to the media to select for GFP positive cells. Clone ID V2LHS_80856 cells did not proliferate. Clone IDs V2LHS_80785 and V2LHS_80786 HuT78 cells were further sorted by the BD FACSaria II cell sorter (BD Biosciences) to obtain >95% pure GFP positive cells. A3G knock-down was verified by measuring the expression of *A3G* by qPCR. While Clone ID V2LHS_80785 did not show any difference in A3G gene expression in the WT and KD cells, clone ID V2LHS_80786 showed a significant reduction in A3G expression and was henceforth used for our studies (KD HuT78 cells) (Fig. 3c).

### RT-PCR and Sanger Sequencing

Total RNA was isolated and reverse transcribed to generate cDNAs as described earlier [15]. DNA primers used for PCR were obtained from Integrated DNA Technologies and are noted in Additional file 7; Table S6. Primers used for PCR of cDNA templates were designed such that the amplicons spanned multiple exons. Agarose gel electrophoresis of PCR products was performed to confirm the generation of a single product in a PCR and then sequenced on the 3130 xL Genetic Analyzer (Life Technologies) at the RPCCC genomic core facility as described previously [17]. To quantify RNA editing level, the major and minor chromatogram peak heights at putative edited nucleotides were quantified with Sequencher 5.0/5.1 software (Gene Codes, MI). Since the software identifies a minor peak only if its height is at least 5% that of the major peak’s, we have considered 0.048 [=5/(100+ 5)] as the detection threshold [17, 27].

For quantitative PCR to assess *APOBEC3G* and *APOBEC3F* gene expression, reactions using LightCycler™ 480 Probes Master and SYBR™ Green I dye were performed on a LightCycler™ 480 System (Roche). Quantification cycle (C_q_) values were calculated by the instrument software using the maximum second derivative method, and the mean C_q_ value of duplicate PCR reactions was used for analysis.

### Immunoblotting assays of cell lysates

Whole cell lysates were prepared and immunoblot was performed as described previously [15, 34]. APOBEC3G antiserum (Apo C17, catalog number-10082) was obtained from the NIH AIDS Reagent program [64, 65], Rabbit monoclonal Phospho-eIF-2α (Ser51) (product number-3398, DG98) was obtained from Cell Signaling Technology, mouse monoclonal anti-β-actin (product number AM4302, AC-15) was obtained from Life Technologies, mouse monoclonal anti-HIF1α (product number GTX628480, GT10211) and rabbit polyclonal anti-α-Tubulin (product number GTX110432) was obtained from GeneTex and used at dilutions recommended by their manufacturers in 5% milk, except Phospho-eiF-2α, which was diluted in 5% BSA. HRP-conjugated goat anti-mouse or anti-rabbit antibodies were purchased from Life Technologies and used at 1:2000 dilution followed by chemiluminescent detection of the proteins [15].

### Cell proliferation assay

WT and KD HuT78 cells (1×10^6^ cells in 100 μl per well) were seeded in 96-well round-bottom plates and incubated covered in the culture medium for 22 hours in a 37°C humidified hypoxia chamber (1% O_2_) or 37°C humidified culture chamber (21% O_2_). Cell viability was determined using a WST-8 viability stain based colorimetric assay (Dojindo Molecular Technologies, Inc.). Plates were read at 450nm on an Epoch2 microplate reader (Biotek) using the Gen5 software (Biotek).

## Statistical Analysis

Statistical analysis was performed using GraphPad Prism (7.03). A3G expression levels and mean editing levels in different cell types (Fig. 1) were first determined to be significantly statistically different by 1-way ANOVA followed by the recommended multiple comparison tests. RNA editing level and cell proliferation differences between WT and KD Hut78 cells for each gene (Fig. 3e and Fig. 5c) were examined by multiple t tests using the Holm-Sidak method, with alpha=0.05. The effect of inhibitors on RNA editing was first determined to be statistically significant by 2-way (Fig. 4b) or 1-way (Fig. 4d) ANOVA followed by, multiple comparisons of the treatment means for day 1 and/or day 2 using the recommended Dunnett’s multiple comparisons test. Respiration to glycolysis ratios (R/G) were calculated using basal respiration value for each well divided by the average glycolysis value of all wells for each experimental group (n=3 for WT and KD HuT78 cells). These ratios were then normalized to the corresponding WT and KD T0 (unstressed cells) ratios within each experimental group, which are set to 1 (Fig. 5b). The comparison of WT and KD HuT78 cells R/G ratios under stress, across all experiments were performed by Mann-Whitney non-parametric test after normalizing the R/G values against the average of WT stress ratio in experiment 1. P values are indicated by stars: *=p<0.05, **= p<0.01, ***= p<0.001, **** =p<0.0001.

## Others

Gene expression analysis of A3G is performed on two online platforms: (1) BIOGPS at http://biogps.org/#goto=welcome, a collection of thousands of gene expression datasets and (2) Cancer Cell Line Encyclopedia (CCLE) portal at https://portals.broadinstitute.org/ccle. CCLE database contains 1457 cell lines. Weblogo is created at http://weblogo.berkeley.edu/ (2/19/18) with default parameters[66].

## Declarations

### Ethics approval

Not applicable.

### Consent for publication

Not applicable.

### Availability of data and materials

The RNASeq data of NK cells have been deposited in the Gene Expression Omnibus (GEO) data bank, accession code GSE114519.

### Author contributions

BEB conceived the study and designed the experiments with contributions from SS. SS performed most of the experiments. BEB and SS wrote the manuscript. ECG and JW analyzed the RNASeq data and wrote the method for the same in the manuscript; other bioinformatics and statistical analysis was performed by BEB. SP performed cell viability assays with support from ESW. OM performed flow cytometry to test the purity of primary NK, CD4+ T and CD8+ T cells. PHB contributed toward performing the experiment with NK-92 cells. All authors read and approved the final manuscript.

## Acknowledgements

Flow Cytometry, RNASeq, Sanger sequencing and lentiviral knockdown of A3G in HuT78 T cells services were provided by the Flow and Image Cytometry, Genomics, and Gene Modulation Services, respectively at RPCCC’s shared resources facility; which are partly supported by NCI Cancer Center Support Grant 5P30 CA016056. The following reagent was obtained through the NIH AIDS Reagent Program, Division of AIDS, NIAID, NIH: anti-ApoC17 from Dr. Klaus Strebel.

